# High-temporal-resolution point-of-care multiplex biomarker monitoring in small animals using microfluidic digital ELISA

**DOI:** 10.1101/2025.05.11.653356

**Authors:** Yujing Song, Andrew D. Stephens, Huiyin Deng, Adrienne D. Füredi, Shiuan-Haur Su, Yuxuan Ye, Yuxiang Chen, Michael Newstead, Qingtian Yin, Jason Lehto, Zeshan Fahim, Benjamin H. Singer, Katsuo Kurabayashi

## Abstract

Time-course monitoring of blood biomarkers with rapid turnaround has the potential to revolutionize the diagnosis, stratification of phenotypes, and therapeutic/prognostic approaches for various acute inflammatory diseases in both clinical and preclinical studies. Current approaches, however, are hampered by slow turnaround times and large sample volume requirements, limiting the exploration of disease mechanisms and therapeutic strategies. Here, we developed a microfluidic digital ELISA platform prototype, combining single-molecule counting with whole blood assay capability for the first time from small animal models. This platform is automated and enables repeated, rapid biomarker monitoring with just 3.5 µL of whole blood collected from the tail. Our platform demonstrated high sensitivity and multiplexity, allowing real-time cytokine profiling within a 2-hour turnaround. Using a murine sepsis model, we achieved precise temporal monitoring of cytokine levels, demonstrating prognostic capability by correlating early-stage cytokine levels with a liver-injury biomarker. This microfluidic platform enables high temporal resolution and rapid monitoring of biomarker dynamics in a single mouse using freshly collected whole blood, significantly reducing the number of animals needed for preclinical studies. This technology has strong potential to transform ICU therapeutic strategies and preclinical research, enabling personalized treatment based on real-time biomarker profiles.

## 1. Introduction

Point-of-care monitoring of biomarkers is crucial for understanding biological processes and disease progression, especially in acute inflammatory conditions such as Sepsis (Ashley and Hassan, 2021; Bradley and Bhalla, 2023; Park et al., 2020; Reddy et al., 2018), ARDS (Maddali et al., 2022), COVID-19 (Q. Song et al., 2021; Suleman et al., 2021; Yang et al., 2021; Yessayan et al., 2020), and cancer immunotherapy (Williamson and Mendes, 2024; Y. Song et al., 2021a). Over the last decade, retrospective analyses of clinical trials that involve detecting biomarkers from banked blood samples have demonstrated the existence of hyperinflammatory and hypoinflammatory subphenotypes in ARDS, pneumonia, and sepsis (Sinha et al., 2024, 2023; Famous et al., 2017). These studies suggest that patients with different subphenotypes respond differently to supportive care. Timely and prospective immunomodulatory therapy, guided by point-of-care subphenotype differentiation, has the potential to significantly improve clinical outcomes (Maddali et al., 2022). The first step in developing such a therapeutic strategy is to establish a diagnostic algorithm based on dynamic biomarker profiles and understand the evolution of inflammatory responses over time in individual patients.

Small animal models are a crucial tool for understanding critical illness syndromes and the relationship of immunopathology to organ injury on an individual level (Matthay et al., 2024). However, elucidating biomarker dynamics in small animals is constrained by the need for large sample volumes and insufficient temporal resolution (Lee et al., 2018). In the U.S., preclinical studies utilize more than 100 million mice annually (Carbone, 2021), leading to high costs and significant ethical concerns. Conventional animal model studies (Pan et al., 2017; Vasalou et al., 2024), particularly those examining immune disorder pathogenesis, often involve sacrificing animals for single-time-point biomarker measurements. This method restricts the ability to comprehensively understand time-evolving immune status during inflammatory conditions. Current immunoassay platforms require blood volumes that are impractical for serial sampling in small animals, necessitating the use of multiple animals across different time points. As such, there is a need for advanced blood biomarker detection methods that offer fast turnaround times, enable multi-time-point blood draws with very small sample volumes, eliminate extensive sample preparation steps, and provide immediate, actionable insights.

This study aims to address the need for improved biomarker monitoring by developing a microfluidic digital ELISA platform for multiplexed, fine-time monitoring of whole blood biomarkers in small animals. Existing microfluidic immunoassay technologies have shown promise in reducing sample volumes and improving sensitivity (Dong and Ueda, 2017; Fan et al., 2008). However, conventional assays often fall short in providing the temporal resolution and sensitivity required for detailed biomarker dynamics in small animals (Lee et al., 2018). Recent advancements in single-molecule counting digital ELISA have significantly improved sensitivity compared to traditional methods (Cohen et al., 2020; Maley et al., 2020; Rissin et al., 2010; Wu et al., 2022), enabling the detection of biomarkers at sub-femtomolar concentrations. This is particularly advantageous for small animal studies, as it permits high dilutions and smaller sample volumes. More recently, Wang et al. developed a microfluidic digital immunoassay for point-of-care detection from whole blood using a plasma separation filter (Chen et al., 2024). Despite these advancements, most existing digital ELISA platforms continue to rely on magnetic beads suspended in-reaction, which necessitates sample preparation and large sample volumes, making the techniques inadequate for rapid and minimally invasive sampling. To the best of our knowledge, no digital ELISA has yet achieved direct whole blood detection without the need for plasma separation, a capability we introduce with our platform.

In previous work (Y. Song et al., 2021c, 2021b; Stephens et al., 2023), we demonstrated the first prototype of a digital immunosensor platform named the “Pre-equilibrium digital ELISA (PEdELISA).” The PEdELISA technology lays the foundation for high-sensitivity, ultrafast biomarker detection by leveraging microfluidics and single-molecule detection. In this study, we have further advanced the PEdELISA platform with a new prototype capable of detecting biomarkers directly from whole blood. This system integrates pre-patterned microarrays within a microfluidic chip featuring internal passivation coatings that prevent cell adhesion and hemolysis, enabling “on-chip” whole blood analysis – a key difference from conventional digital ELISA. Additionally, we have achieved the automation of the platform, allowing it to function as a stand-alone system suitable for deployment in point-of-care testing at a clinical or bio-research lab. This automated prototype streamlines the workflow, reducing human error resulting from manual operations, thereby enhancing the platform’s reliability and ease of use.

Our research demonstrates the platform’s capability to achieve 2-hour temporal resolution of cytokine profiling on a single mouse at the point-of-care from blood draw to data delivery, using only one drop of whole blood (3.5 μL) at each time point. This advancement has facilitated fine-time and point-of-care monitoring of pro-inflammatory chemo/cytokines in cecal slurry (CS) mouse models, providing detailed temporal profiles associated with the systemic inflammatory response to various sepsis-induced conditions that mimic complex bacterial infections, anti-inflammatory treatment, and endpoint liver injury.

## 2. Materials and methods

### 2.1. Materials

Mouse IL-6 and MCP-1 capture and biotinylated detection antibody pairs, ELISA MAX™ Deluxe Set, ELISA Assay Diluent B, and HRP Streptavidin were purchased from BioLegend. Mouse CXCL-1 and CCL-11 capture and biotinylated detection antibody pairs, and DuoSet ELISA kits were obtained from R&D Systems. Dynabeads™ M-270 Epoxy beads, QuantaRed™ Enhanced Chemifluorescent HRP Substrate Kit, SuperBlock™ (PBS) Blocking Buffer, and Blocker™ Casein were obtained from Thermo Fisher Scientific. Sylgard™ 184 PDMS was purchased from Dow Corning. Novec™ 1720 fluorosilane polymer and Novec™ 7500 fluorinated oil were purchased from 3M™, and 100 mm P-type silicon wafers were from University Wafer Inc. (Boston, MA).

### 2.2. Ethics Statement

The Institutional Animal Care and Use Committee of the University of Michigan approved all animal studies (PRO00010712). The University of Michigan’s laboratory animal care policies follow the Public Health Service Policy on Humane Care and Use of Laboratory Animals.

### 2.3. Animals

Eight- to ten-week-old male C57BL/6 mice were obtained from Jackson Laboratories (Bar Harbor, ME, USA). Mice were housed at 21 °C with a 12-hour light-dark cycle. They had ad libitum access to water and chow (Inotiv Teklad, West Lafayette, IN, USA), and a humane endpoints policy was followed. Mice were weighed both pre-infection and at the time of harvest. Rectal temperatures were obtained at the same time points. Mice were euthanized by CO2 inhalation.

### 2.4. Cecal Slurry Derivation

Cecal slurry was harvested from mice obtained from Jackson Laboratories months apart to increase microbiome variability, following the method of Starr and colleagues (Starr et al., 2014). Mice (n=30) were sacrificed within one week of arrival from Jackson Laboratories, and the ceca was removed.

Cecal contents were collected using flame-sterilized forceps and a spatula, then combined and weighed. Cecal contents were mixed with sterile water at a ratio of 0.5 mL water to 100 mg cecal contents. This mixture was sequentially filtered through 810 μm and 20 μm sterile screens and mixed with an equal volume of 30% glycerol in PBS. This slurry was vigorously mixed, aliquoted, and stored at −80°C until use.

### 2.5. Infection with Cecal Slurry

To induce sepsis, 14-18 μL/g of cecal slurry stock per gram of body weight was injected intraperitoneally. Control animals were injected with an equal volume of saline according to body weight. Mice received ceftriaxone (75 mg/kg) and metronidazole (25 mg/kg) via intraperitoneal injection and 1 mL of warmed saline via subcutaneous injection 6 hours after infection. Infection was monitored with rectal temperature measurements.

### 2.6. PEdELISA Chip Assembly and Surface Function

See Supplementary Figure S1 for chip fabrication and surface function details.

### 2.7. Programmed PEdELISA Assay

The assay began with priming the manifold with the washing buffer (WB: PBS-T 0.1% Tween20). WB was loaded into the inlet reservoirs and then extracted across the flow cell driven by the negative pressure from the syringe pump. Diluted whole blood samples (35 μL) were then added to each sample reservoir and similarly extracted across the flow cell by the pump. The samples were allowed to react with the sensors for 10 minutes, followed by quenching the reaction by flushing the flow cells with 100 μL of WB, and then performing a continuous flushing with a total of 5 mL of WB for around 5 minutes. Next, detection antibody cocktails were introduced and allowed to react in all flow cells for 5 minutes, followed by a continuous flushing with 3 mL of WB. The system then drew in 40 μL of the avidin-HRP solution (100 pM) and gradually loaded it into the chip for enzyme labeling, which took 3 minutes. The chip was washed again with a total of 5 mL of WB for 5 minutes. Finally, the system drew and loaded 25 μL of the QuantaRed (Qred) substrate solution, sealed it with 50 μL of fluorinated oil (HFE-7500, 3M), and initiated the image scanning process.

### 2.8. Whole blood and Plasma Correlation Assay

Ten mice were injected with 18 μL/g of cecal slurry (CS), while two uninjected mice served as negative controls. At the 2hr time point, 4 infected mice and 2 uninfected mice were euthanized and sacrificed, with the remaining 6 mice euthanized at the 5hr. Central blood samples were collected via syringe at each time point and immediately transferred into two anticoagulant buffers: 100 μL of blood into 900 μL of heparinized saline (25 U/mL) or 900 μL of EDTA saline (5 mM). The samples were then split into two aliquots (500 μL each): one for whole blood analysis and the other processed to obtain diluted plasma by centrifugation (2000 rcf for 15 min at 4°C). All samples were stored on ice and measured by PEdELISA within 12 hours. Prior to the PEdELISA assay, samples were gently mixed by inversion and further diluted 30-fold in ELISA dilution buffer (1x PBS, 1% BSA).

### 2.9. Blood Sampling

For all cytokine measurements, blood was drawn from a nick in the tail vein using a heparinized glass capillary tube pre-marked at a 3.5 μL volume (Micro-Cal tube, Kimble-Chase). The sample was immediately ejected using a pipette into a 100 μL volume of heparinized (25 U/mL) saline. At the time of euthanasia, blood was drawn from the right ventricle using a syringe and inoculated into a low-volume serum separator tube (Microvette, Sarstedt).

### 2.10. AST Assay

Whole blood was collected into serum separator tubes, allowed to clot, and separated into serum by centrifugation. Serum chemistries were run on a Liasys 330 (AMS Alliance, Guidonia, Italy) automated wet chemistry analyzer. Assays were performed within the ULAM In Vivo Animal Core pathology laboratory at the University of Michigan. Quality control was performed daily using manufacturer-provided reagents. The laboratory is a participant in an external independent quarterly quality assurance program (Veterinary Laboratory Association Quality Assurance Program).

### 2.11. Statistical Analysis

All results are expressed as the means ± standard error of the mean (SEM). GraphPad Prism was used for data analysis. Pearson’s R² value was used to quantify the PEdELISA to ELISA correlation and the correlation between cytokine values and liver injury, by comparing cytokine values at each time point with the value of AST at 24 hours post-infection.

## 3. Results and discussion

### 3.1. Concept of PEdELISA-enabled longitudinal mouse model study

In this study, we consider the evolution of circulating cytokine expression in the murine model of sepsis induced by cecal slurry (CS) treated with antibiotics 6 hours after sepsis induction (Bongers et al., 2024; Starr et al., 2014). Conventional methods for studying immune activation over time in sepsis models typically requires sacrificing the animal at each time point to collect a total blood volume of 1–2 mL and extracting serum or plasma for analysis. This approach necessitates multiple animals for multi-time-point analysis (Figure 1A) and can produce confounding results due to variations from both changes in the host’s immune status over time and host-to-host heterogeneity. Achieving high temporal resolution with conventional methods is prohibitively resource-demanding and expensive due to the need for a large number of animals. Additionally, this approach makes it impossible to correlate early immune events with late organ injury on an individual, within-subject basis.

**Figure 1.**
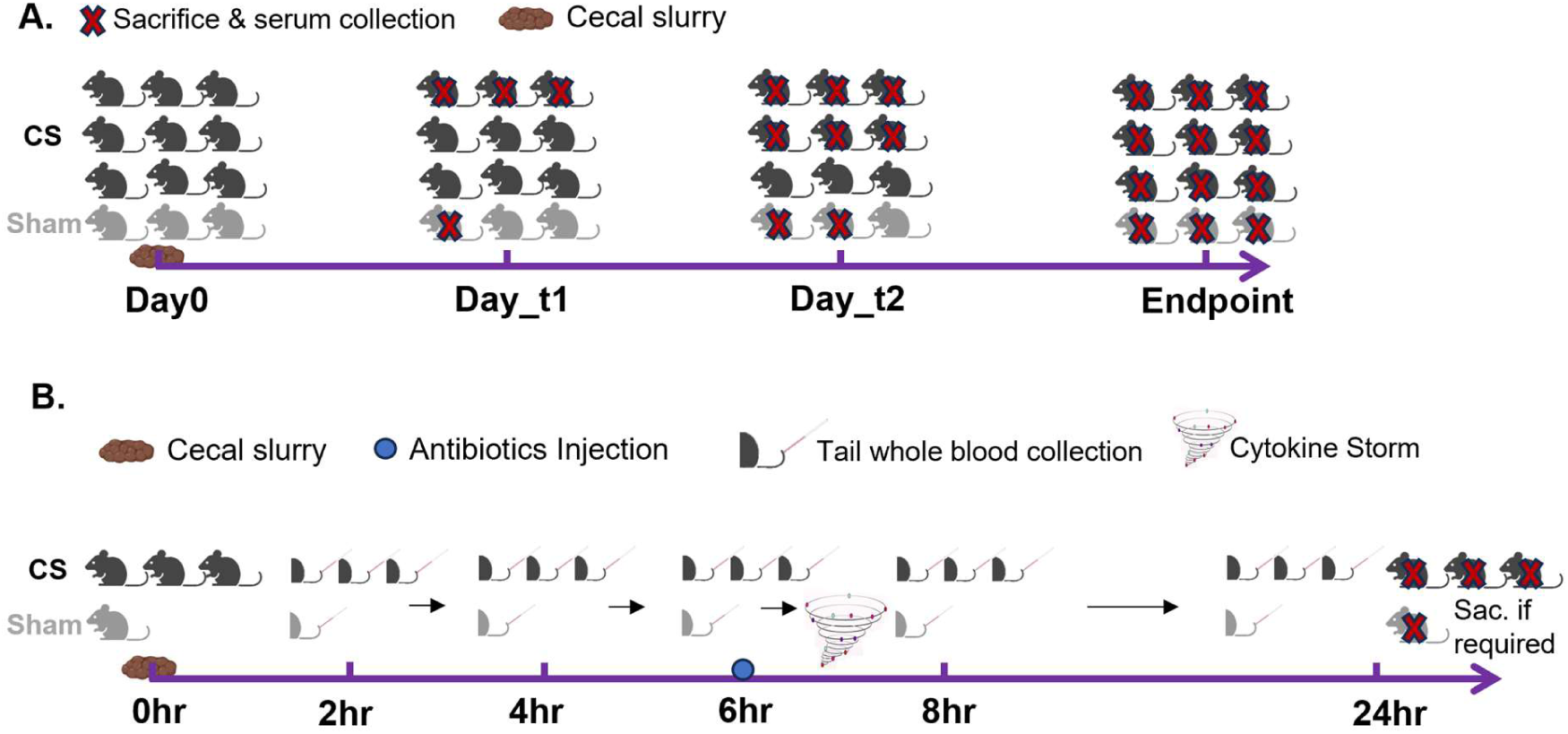
Concept of PEdELISA-enabled time-course mouse model studies. (A) Schematic of conventional time-course mouse model study requiring animal sacrifice and serum collection at a single time point, where each time point contains 3 infected and 1 sham mouse. This method is retrospective, with daily time resolution, large sample volume requirements, and high costs due to the number of animals needed. (B) Schematic of the new mouse model study enabled by PEdELISA, which eliminates the need for animal sacrifice at each time point. This approach allows real-time, prospective monitoring with hourly time resolution using tail whole blood collection (3.5μL), significantly reducing costs and the number of animals required, and enabling the collection of within-subject data over time.

In contrast, the PEdELISA-enabled longitudinal sampling study allows for multi-time-point monitoring of blood biomarkers while keeping the animal alive, even during illness (Figure 1B). This approach enables observation of how the specific mouse’s inflammatory condition evolves over time, as indicated by the temporal biomarker profiles. In this study, 3.5 μL of tail whole blood was collected at multiple time points using a marked capillary tube, diluted 10 times in heparinized normal saline solution, and immediately prepared for the PEdELISA assay (see Materials and Methods). This method provides up to hourly time resolution with minimal blood volume per collection, eliminating extensive sample preparation, significantly reducing costs, and decreasing the number of animals required.

### 3.2. PEdELISA Engineering Prototype

We developed a new engineering prototype of the PEdELISA system with automation, allowing it to function as a stand-alone system at the point of care. Figure 2A provides a schematic and image of the newly designed PEdELISA chip, which integrates a low-cost, soft lithography-molded microwell structures on the bottom substrate with a flow cell layer made of laser-cut polymethyl methacrylate (PMMA) and pressure-sensitive adhesives (PSA) (Figure S1). Unlike conventional digital immunoassays (Wilson et al., 2016), which typically require off-chip in-suspension reactions and a pipetting workstation for automated sample/reagent handling, the PEdELISA system confines antibody-conjugated beads into femtoliter-sized microwells arranged in a pre-patterned array. Each well forms a fluorescent “pixel” upon target analyte binding, enabling the entire assay workflow to be performed “on-chip.” This design facilitates whole blood assays with a minimal sample volume, while preventing hemolysis and bead aggregation during sample processing.

**Figure 2.**
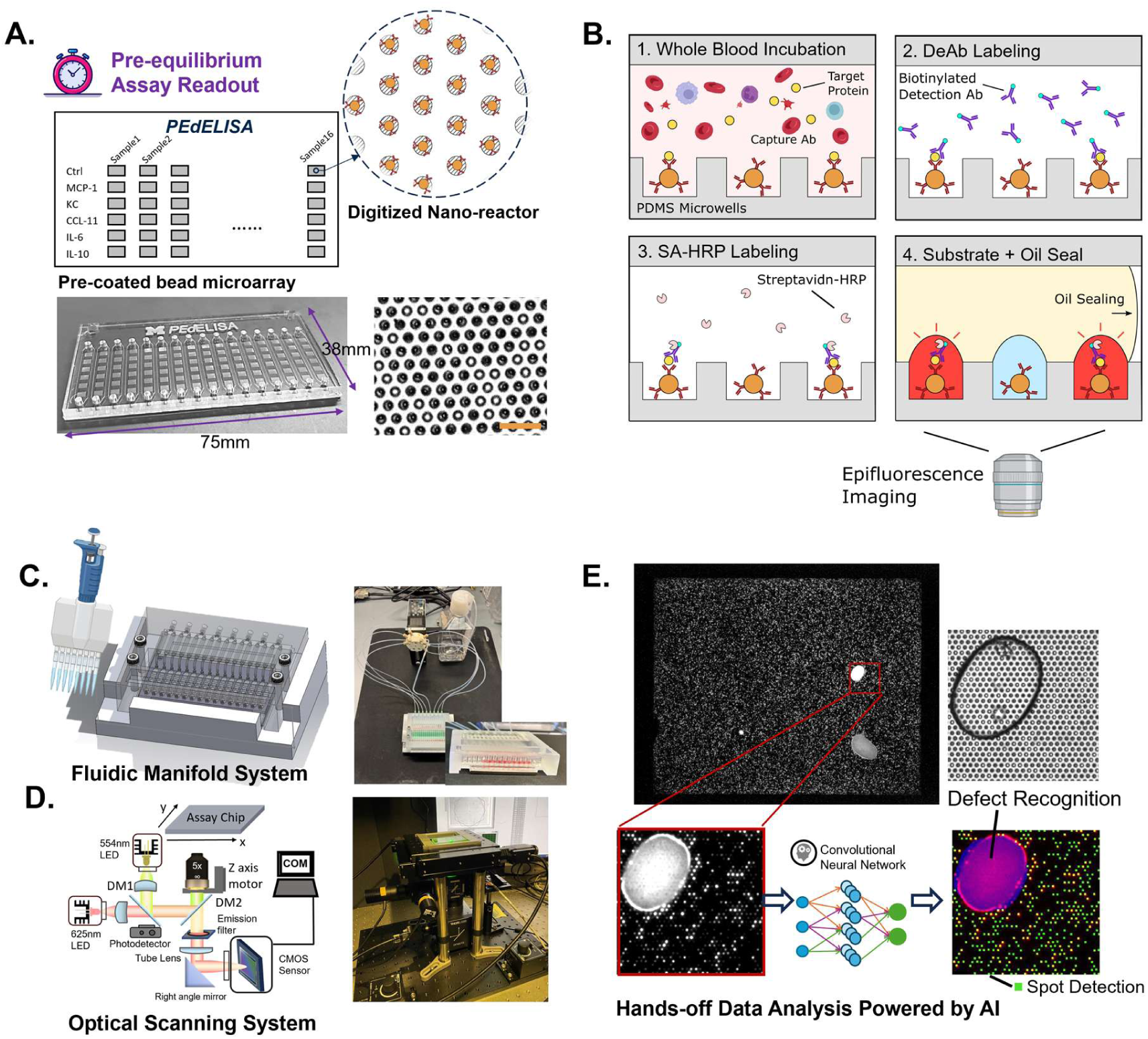
Engineering Prototype of PEdELISA. (A) Chip design concept and image of the PEdELISA chip, illustrating the pre-equilibrium concept and pre-patterned microarray technology for multiplex biomarker detection. Scale bar: 20 µm. (B) Current workflow of the PEdELISA assay, which includes whole blood incubation, detection antibody labeling, streptavidin-HRP labeling, substrate loading, oil sealing, and fluorescence imaging. (C) Design and real image of the PEdELISA fluidic system (inset: manifold with whole blood incubation). (D) PEdELISA reader system and graphic user interface (GUI) for high-throughput, fully-automated fluorescence imaging. (E) Automated data analysis powered by AI. Fluorescence and brightfield images obtained from the PEdELISA assay were analyzed by a pre-trained convolutional neural network (CNN).

The chip is designed to support up to 6-plex detection of biomarkers, although higher plexity measurements are possible (Y. Song et al., 2021c). For demonstration, we chose to detect a panel of MCP-1, CXCL-1, CCL-11, and IL-6, allowing simultaneous detection of 16 different samples on-chip with the current flow cell design (Figure 2A). IL-6 is a common marker of pro-inflammatory phenotypes, and MCP-1 and CXCL-1 are important monocyte and neutrophil chemoattractants, respectively (Sinha et al., 2023). CCL-11 is a marker of inflammation and brain dysfunction in aging (Villeda et al., 2011). The PEdELISA assay workflow includes whole blood incubation, detection antibody labeling, streptavidin HRP labeling, substrate loading, oil sealing, and fluorescence imaging (Figure 2B). Automated reagent manipulation takes 30-40 minutes, with imaging requiring less than 5 minutes. Including sample handling, loading, and analysis, the operational assay time is around 1 hour for up to 16 samples and 6-plex detection.

To further enhance usability, we developed a semi-automated fluidic system with a user-friendly graphic user interface (GUI) for the PEdELISA platform, as shown in Figure 2C. This system consists of the PEdELISA chip, a 3D-printed fluidic manifold, a syringe pump, and a waste bottle. Users load 35μL of the diluted whole blood using a multi-channel pipette, while the GUI controls the programmable syringe pump and manifold system to automatically drive the liquids into the PEdELISA chip, performing the assay and washing steps (see Materials and Methods). To simplify the data acquisition process, we also constructed a compact, cost-effective PEdELISA reader system (Figure 2D). This system features a GUI for user input setup, quick scanning, and auto-focusing of the chip using a custom machine vision algorithm (Figures S2-3). Brightfield/fluorescence image acquisition based on user settings is completed within 5 minutes.

Figure 2E shows the fluorescence and brightfield images obtained from the PEdELISA assay. These images are processed using a custom-trained convolutional neural network (CNN) (Gao et al., 2021; Y. Song et al., 2021c), which performs fluorescence spot detection and counting, identifies image dust and defects, counts the brightfield bead filling rate, and calculates the digital immunoassay signal as “Average Enzyme Molecule per Bead (AEB).” The CNN algorithm facilitates hands-off data analysis powered by AI.

### 3.3. PEdELISA whole blood assay characterization

To evaluate the performance of the newly developed PEdELISA engineering prototype, we conducted experiments to characterize the assay’s sensitivity, specificity for multiplex detection, accuracy, and whole blood assay capabilities. Figure 3A presents the titration standard curves for four biomarkers. Recombinant proteins were mixed at final concentrations ranging from 10 ng/mL to 1 pg/mL and measured using multiplexed PEdELISA. Each data point represents the average signal from three independent chips, with data fitted using four-parameter logistic (4-PL) regression to calculate the limit of detection (LOD).

**Figure 3.**
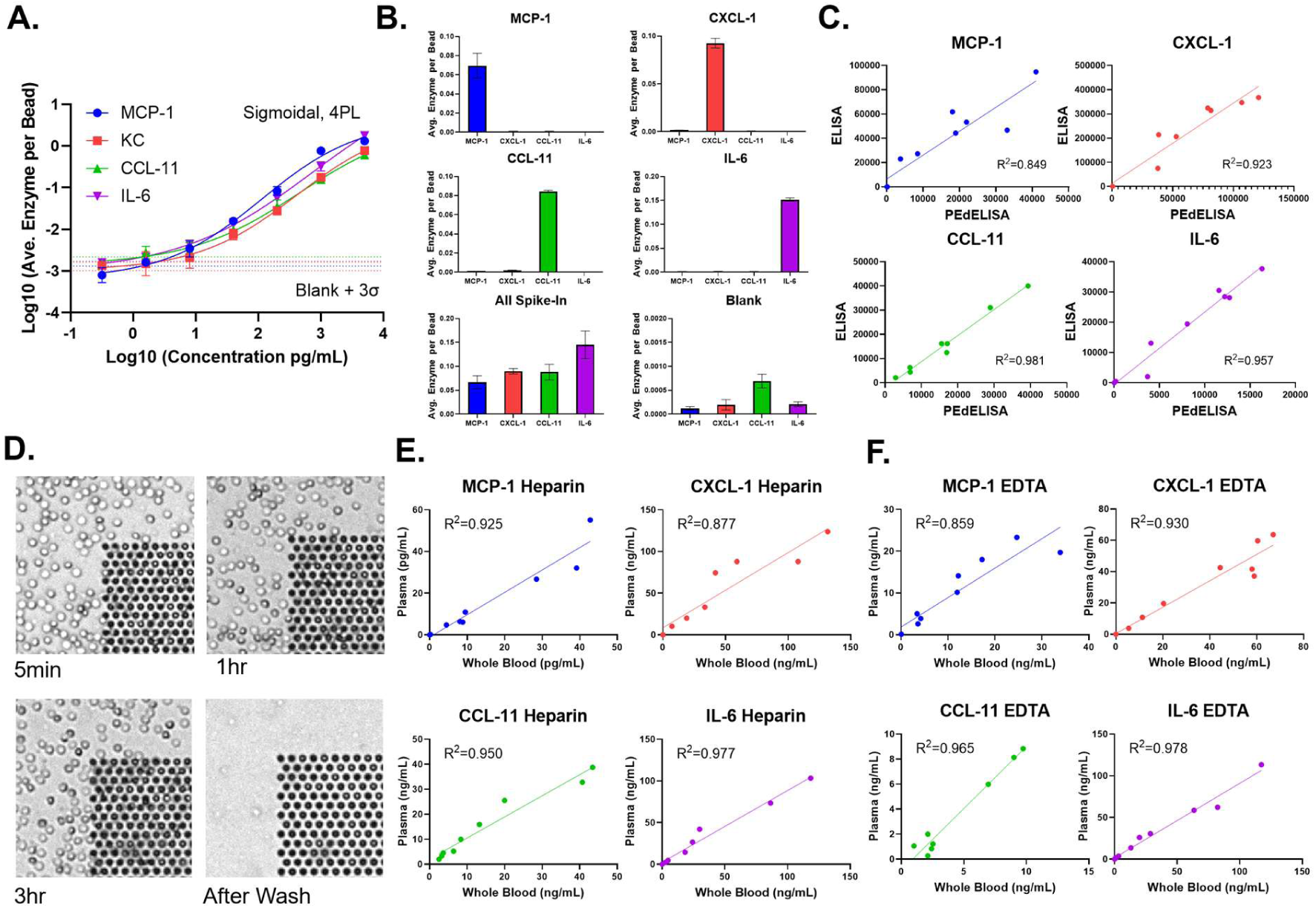
Whole Blood Assay Characterization. (A) 4-plex PEdELISA titration standard curves from 0.32 pg/mL to 5000 pg/mL in ELISA dilution buffer (1% bovine serum albumin). Each data point represents the average signal from three independent chips. The digital immunoassay signal (average enzyme per bead, AEB) was fitted with four-parameter logistic (4PL) curves. The dotted line represents the signal level from a blank solution plus 3 times the standard deviation (3σ) for each cytokine, which is used to estimate the limit of detection (LOD). (B) Assay specificity test with “all-spike-in,” “single-spike-in,” and “blank” (negative) samples of recombinant cytokine marker(s) at 400 pg/mL in ELISA dilution buffer. (C) Correlation between multiplex PEdELISA and conventional single-plex ELISA results using diluted mouse plasma from tail blood collection. Pearson’s R² values were calculated for MCP-1 (R^2^=0.849), CXCL-1 (R^2^=0.923), CCL-11 (R^2^=0.981) and IL-6 (R^2^=0.957). (D) On-chip blood culture test at 5min, 1hr, 3hr and after wash, validating chip surface passivation. Mouse whole blood was diluted 10-fold in 1x PBS. (E) Correlation between whole blood and plasma using Heparin as the anticoagulant measured by PEdELISA using freshly collected CS mouse samples. Pearson’s R² values were calculated for MCP-1 (R^2^=0.925), CXCL-1 (R^2^=0.877), CCL-11 (R^2^=0.950) and IL-6 (R^2^=0.977). (F) Correlation between whole blood and plasma using EDTA as the anticoagulant measured by PEdELISA using freshly collected CS mouse samples. Pearson’s R² values were calculated for MCP-1 (R^2^=0.859), CXCL-1 (R^2^=0.930), CCL-11 (R^2^=0.965) and IL-6 (R^2^=0.978).

To assess the assay’s specificity, we spiked each recombinant cytokine or chemokine analyte into an ELISA dilution buffer (1% BSA) to simulate a high-protein background matrix. Figure 3B shows the results for “all-spike-in,” “single-spike-in,” and “no-spike-in” samples using 400 pg/mL recombinant cytokine markers, demonstrating negligible antibody cross-reactivity between analytes. For accuracy validation, we compared the multiplex PEdELISA results with the gold standard single-plex ELISA methods by retrospectively measuring banked mouse plasma samples for each cytokine. Figure 3C displays the correlation between PEdELISA and ELISA, with typical R-square values of 0.85-0.98 for the biomarkers.

To enable whole blood detection, we developed a surface treatment protocol using Novec™ 1720 fluorosilane polymer (3M) followed by SuperBlock™ (ThermoFisher) passivation to prevent cell adhesion and minimize hemolysis (see Materials and Methods). We evaluated the performance of this protocol by imaging PEdELISA flow cells during and after exposure to a representative mouse whole blood sample. Figure 3D shows brightfield images of 10x diluted whole blood incubated in the PEdELISA flowcells at 5 minutes, 1 hour, and 3 hours, followed by a wash step to remove residual blood cells. No significant red blood cell lysis or cell adhesion was observed during the 3-hour incubation, which greatly exceeds the typical 10-min whole blood assay incubation time.

In addition to freshly drawn whole blood, the PEdELISA system is compatible with various sample types, as demonstrated in our previous work with serum, plasma, and cell culture medium (Su et al., 2021, 2023). Here, we conducted a controlled comparison of heparin and EDTA whole blood with their corresponding diluted plasma using samples from a CS sepsis mouse model. Blood was collected at 2 and 5 hours post-infection, diluted 1:10 in heparinized (25 U/mL) or EDTA saline (5 mM), and then split into two aliquots: one for whole blood analysis and the other processed to obtain diluted plasma (see Materials and Methods). All samples were stored on ice and measured by PEdELISA within 12 hours. We observed consistent correlations between whole blood and diluted plasma for both anticoagulant types, with R² values ranging from 0.88 to 0.98 for heparin, and 0.86 to 0.98 for EDTA (Figure 3E,F), which validates PEdELISA’s capability for accurate whole blood detection.

### 3.4. PEdELISA high-temporal-resolution whole blood biomarker detection

To demonstrate the unique features of the PEdELISA system, we collected and stored small amounts of blood (3.5 μL) from the tail vein every hour (or every 2 hours, depending on the mouse’s CS dose) for up to 12 hours. Within 24 hours of sample collection, we loaded the 12 stored blood samples for the same mouse—representing different collection times—onto the PEdELISA chip to measure four chemo/cytokine markers simultaneously. Figure 4A presents the high-resolution biomarker profiles measured for two mice injected with different doses of CS contents: Mild (14 μL/g), Severe (18 μL/g), and two control mice injected with normal saline, all treated with antibiotics (ceftriaxone and metronidazole) 6 hours after infection. Rectal (body) temperature was measured every hour for all mice across the three conditions.

**Figure 4.**
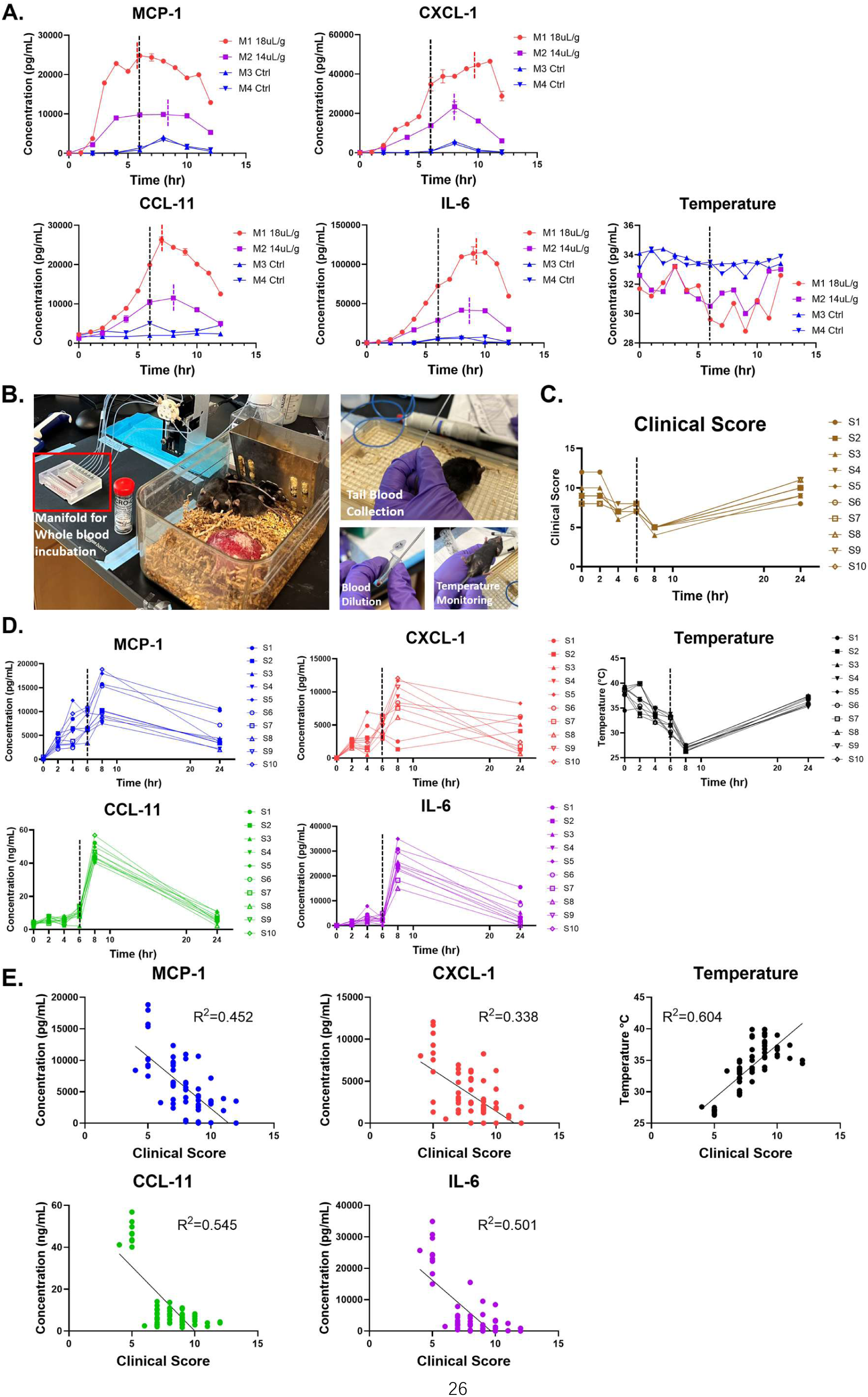
PEdELISA high-temporal-resolution whole blood biomarker detection. (A) High-resolution retrospective experiment to evaluate CS-induced sepsis severity, disease development dynamics, and response to antibiotics at 6 hours post-injection. Whole blood samples were collected every hour for the high dose case (18uL/g) and every 2 hours for the low dose (14uL/g) and control (saline) cases for a total of 12 hours. Rectal (body) temperature was monitored every hour. Antibiotics (ceftriaxone and metronidazole) were administered at the 6-hour after infection. (B) Photographs of the animal and equipment setup. (i) A cage housing the test mice was placed next to the manifold operation unit for the PEdELISA chip. (ii) Whole blood was drawn from the tail vein of one of the mice. (iii) The collected blood sample was diluted by a buffer solution within the PCR tube and subsequently loaded into the inlets of the manifold for PEdELISA biomarker analysis on the microfluidic chip. (iv) The rectal temperature of the mouse was measured. (C) Clinical scores recorded for the mice based on their response to a finger poke, signs of encephalopathy, and overall appearance, evaluated every 2 hours. (D) Real-time 4-plex whole blood detection results of 10 mice in parallel using PEdELISA, showing cytokine levels and body temperature trends over time. 3.5uL of whole blood samples were collected from the tail vein every 2 hours at 0, 2, 4, 6, 8, and 24 hours, diluted, and assayed immediately within a 2-hour turnaround time, showcasing the prospective diagnostic detection capability. Antibiotics were administered at the 6-hour after infection. (E) Correlation between the 4 cytokines, temperature, and clinical scores of the 10 mice at all time points. Pearson’s R² values were calculated and used to determine the correlation quality.

Here, we observed distinct cytokine concentration progressions for different markers. For example, MCP-1 exhibited a rapid increase in response to CS-induced sepsis at the 3-hour time point and remained at a similar level through the later phase, while the levels of CXCL-1, IL-6, and CCL-11 peaked around the 7-9 hour time point, synchronizing with the maximum level of septic shock manifested by the largest body temperature drop. Various cytokines also responded differently to antibiotic treatment, with MCP-1 and CCL-11 levels beginning to decrease two hours after administration, while CXCL-1 and IL-6 levels continued to increase for another four hours. The progression of cytokine concentrations upon antibiotic treatment also varied depending on the CS doses, with several cytokines continuing to increase for several hours in higher dose (18 μL/g), while in the lower dose (14 μL/g), the cytokine storms seemed to be controlled immediately, resulting in reduced cytokine level. Data for the mice in the control group confirmed that this blood collection method did not cause harm or induce a significant inflammatory response in the animals, even with high-frequency blood sampling. A small increase in MCP-1 and CXCL-1 was observed in the control group at the 8-hour time point, which might be due to the injection of normal saline at 6 hours.

Our data also revealed that cytokine monitoring offered a valuable complement to body temperature monitoring for stratifying septic conditions in small animals. While CS mice received higher dose and lower dose exhibited similar body temperature drops as sepsis progressed, their overall inflammatory cytokines were different (Figure 4A). Timely cytokine measurements provided insights into specific immune mechanisms and potential targets for intervention, beyond the global biomarkers indicated by temperature readings.

### 3.5. Point-of-care PEdELISA longitudinal whole blood biomarker monitoring

Subsequently, we demonstrated point-of-care longitudinal measurement for each of 10 mice receiving 14 µL/g CS. Blood was drawn at each time point and immediately analyzed using the PEdELISA system, allowing for real-time biomarker analysis enabled by the engineering prototype (Figure 4B). We collected longitudinal biomarker data every 2 hours for up to 8 hours, with endpoint data collected at 24 hours. The gap between 8 hours and 24 hours in measurements was due to logistical constraints, as continuous blood draw and analysis next to the mouse were not logistically feasible for more than 8 hours. Additionally, we recorded clinical scores for the mice based on their response to a finger poke, signs of encephalopathy, and overall appearance. These scores were used to assess each mouse’s clinical symptoms and illness progression every 2 hours (Figure 4C, see Materials and Methods).

Figure 4D shows the results of point-of-care measurements for 10 mice monitored in parallel. A moderate dose of CS (14 µL/g) was administered for sepsis induction at 0 hour, and 3.5 μL of tail blood was collected at 0, 2, 4, 6, 8, and 24 hours, diluted, and immediately measured using the PEdELISA system. Antibiotics were administered at the 6-hour time point. The data revealed a similar trend for pro-inflammatory cytokines compared to the retrospective study, with notable variability in each animal’s response to bacterial infection. The body temperature trends mirrored cytokine levels, reaching a minimum at 8 hours, with cytokine levels beginning to return to baseline by 24 hours. Figure 4E illustrates the correlation between the four cytokine markers and temperature with the clinical scores of the 10 mice at all time points. Strong correlations between CCL-11, IL-6, and temperature with the clinical score, as evidenced by Pearson’s R² values exceeding 0.5, further highlight the importance of cytokine measurements in monitoring disease progression in sepsis. Table S1 summarizes the detailed clinical scores recorded for the 10 mice.

### 3.6. Association between cytokine levels and liver injury

A disadvantage of cross-sectional study designs that only examine a single time point in each animal is that biological causation requires time lags between an immunopathological event and eventual organ dysfunction. If biomarkers of inflammation and organ injury are measured simultaneously, these time-varying associations are lost. To demonstrate this, we examined the association between cytokine levels, temperature, weight over the course of sepsis and liver injury at 24 hours after sepsis induction. For example, for IL-6, the linear relationship between AST and cytokine levels was strongest at 8 hours post-infection (Figure 5A, Figure S4-5 for other cytokines and R² value). We found that the strength of correlation across most measured cytokines, as measured F-statistic for significance of the linear regression comparing cytokine values to the AST value at 24 hours (Figure 5B), was greatest for the cytokine measurements at 8 hours after CS injection (F(1,23), n=24 mice). In contrast, the strength of temperature to AST was greatest at 24 hours. These time-varying associations underscore the importance of time lags in immunopathology and organ injury. Importantly, some cytokine levels at 24 hours post-infection correlate poorly with other time points (Table S2). For example, for 8hr vs 24hr, the Pearson’s R² are CXCL-1: 0.180 IL-6: 0.303 and cannot be taken as a surrogate for the individual history of inflammation within a single subject.

**Figure 5.**
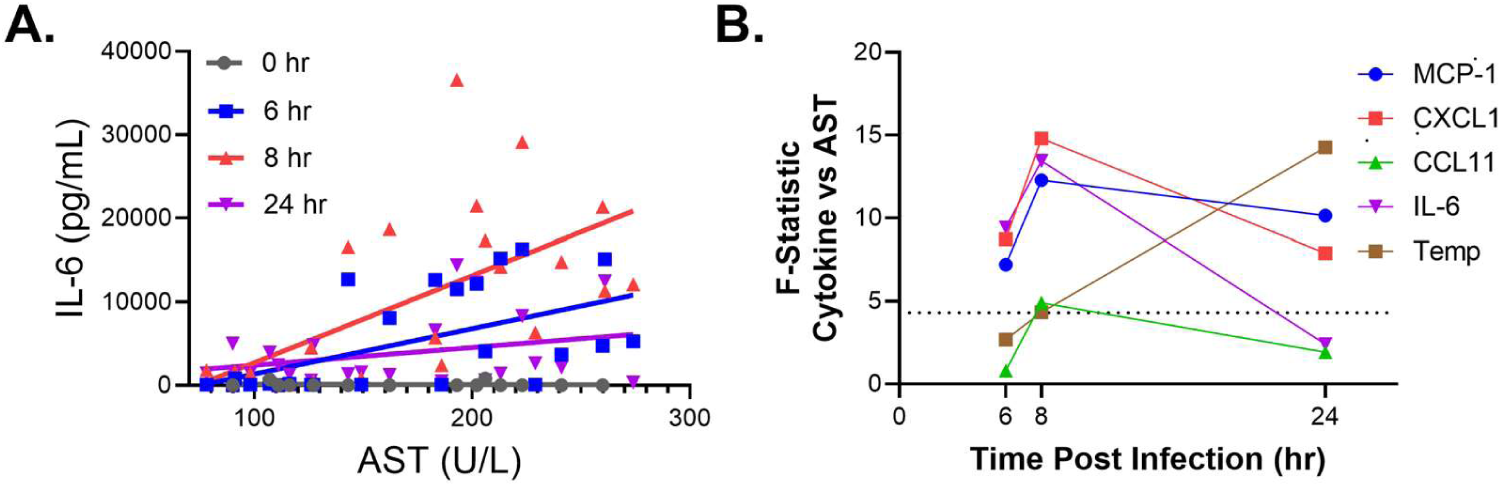
Association Between Cytokine Levels and Liver Injury. (A) Correlation between IL-6 levels at 0, 6, 8, 24 hours post-infection and liver injury marker AST at 24 hours. Cytokine levels measured at 8 hours post infection had the strongest association with liver injury 24 hours post infection. (B) F-statistic values of all cytokines at different times post-infection, correlated with the liver injury marker AST at 24 hours, as measured with the PEdELISA platform, illustrating the importance of time lags in the relationship between immunopathology and organ injury. The dotted line indicates the F(1,23) value corresponding to p = 0.05. n=24 mice.

## Conclusion and outlook

In this study, we introduced the first digital immunoassay capable of direct whole blood detection, demonstrating the PEdELISA platform’s ability to achieve high-temporal-resolution cytokine monitoring in small animal models. This new approach addresses the limitations of conventional methods that require large sample volumes, extensive sample preparation, and often necessitate sacrificing animals for one-time-point measurements. From an engineering standpoint, PEdELISA’s technological breakthrough lies in its whole blood assay capability, uniquely enabled by “on-chip” biosensing, microfluidics, optimized coatings, and microarray patterning. Its compact, automated, stand-alone design ensures practical deployment at the point-of-care. From a biomedical perspective, our findings from the mouse sepsis model highlight the critical need for high-temporal-resolution, minimally invasive whole blood detection to monitor systemic inflammatory disorders. Given the heterogeneity of individual responses and the rapid fluctuations in biomarker levels, PEdELISA’s ability to use just 3.5 μL of blood for multiplexed profiling with a 2-hour temporal resolution provides immediate and actionable insights, with transformative potential for monitoring disease progression and treatment response in both preclinical and clinical settings.

While the PEdELISA platform offers significant advantages, some limitations remain. The current prototype requires manual sample loading, and further work is needed to fully automate the process to reduce human intervention and streamline the workflow. Additionally, current chip manufacturing relies on manual PDMS-molding, patterning, and assembly, which is low-throughput and labor-intensive. Future efforts should focus on scalable manufacturing, such as injection molding, with dry preservation to enable wider use. Expanding biomarker detection capabilities and validating the platform in other disease models will also be critical for broader adoption. Overall, PEdELISA represents a significant advancement in biomarker monitoring, offering the ability to monitor or measure biomarkers from small volumes of whole blood at short intervals of time. If applied to clinical diagnostics or research, this technology has the potential to enable prospective biomarker measurement and provide prognostic insights, making it highly useful for treating patients in critical conditions and for scientific studies.

## CRediT authorship contribution statement

Yujing Song: Conceptualization, Investigation, Methodology, Software, Data curation, Formal analysis, Writing – original draft. Andrew D. Stephens: Investigation, Data curation, Formal analysis, Writing – review and editing. Huiyin Deng: Investigation. Adrienne D. Füredi: Investigation. Shiuan-Haur Su: Investigation. Yuxuan Ye: Investigation. Kevin Chen: Investigation. Michael Newstead: Validation. Qingtian Yin: Investigation. Jason Lehto: Investigation. Zeshan Fahim: Investigation.

Benjamin H. Singer: Conceptualization, Methodology, Funding acquisition, Writing review & editing, Supervision. Katsuo Kurabayashi: Conceptualization, Funding acquisition, Writing – review & editing, Supervision.

## Data availability

Data will be made available on request.

## Acknowledgements

This study was supported by the National Science Foundation (ECCS 1708706 and CBET 1931905, K.K.), the University of Michigan Precision Health Scholars Grant (Y.S.) National Institute of Health (NIH) R01AG074968 and R33HL154249 to B.H.S, and T32HL007749 to A.D.S. H.D. was partially supported by the China Scholarship Council. Device fabrication was performed at the University of Michigan Robert H. Lurie Nanofabrication Facility.

We express our gratitude to Tao Cai, Sonnet Xu, Sachin Agrawal for their contribution to the software development during their summer internships at the University of Michigan.

## Abbreviations

PEdELISA: Pre-equilibrium digital enzyme linked immunosorbent assay; ELISA: Enzyme linked immunosorbent assay; CS: Cecal slury; CNN: Convolutional neural network; GUI: graphic user interface; AEB: Average enzyme molecule per bead; LOD: Limit of detection; PBS: Phosphate buffered saline; BSA: Bovine serum albumin; PDMS: Polydimethylsiloxane; PMMA: Polymethyl methacrylate; PSA: Pressure-sensitive adhesives; MCP-1: Monocyte chemoattractant protein-1; CXCL-1: C-X-C motif chemokine ligand 1; CCL-11: C-C Motif Chemokine Ligand 11, eotaxin; IL-6: Interleukin 6; AST: Aspartate aminotransferase.

## Supplementary data

Supplementary figures and tables are available.

