## Supplementary for "High-temporal-resolution point-of-care multiplex biomarker monitoring in small animals using microfluidic digital ELISA"

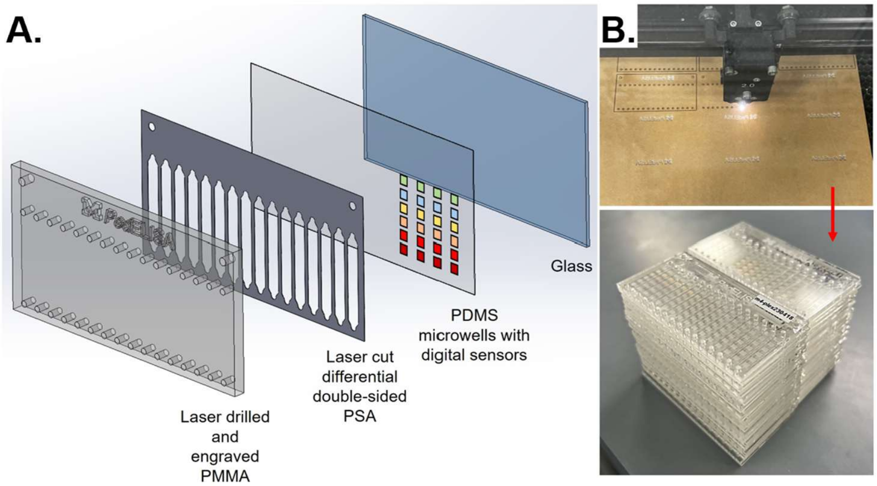


**Figure S1. The architecture and manufacturing of the PEdELISA chip.**

**Fabrication of the PEdELISA Chip**

The PEdELISA chip is fabricated using laser cutting and laminating with pressure-sensitive adhesives (PSA), creating a PMMA-based multilayer structure suitable for mass production (Figure S1). The top layer of the chip, made from 3.175 mm (1/8 inch) cast PMMA, serves as a rigid interface with the fluidic manifold and external pumps. This layer is laser-cut to create 32 inlet and outlet ports and alignment features for interfacing with the manifold. A differential PSA layer with silicon adhesive on one side and acrylic adhesive on the other is laser-cut to form individual flowcells when sandwiched between the PMMA layer and the bottom digital sensor chip. A 3D-printed jig ensures precise alignment and assembly of the PMMA and PSA layers. The assembled flow cells are treated with Novec™ 1720 fluorosilane polymer (3M). The bottom digital sensor component consists of femtoliter-sized microwell arrays made by molded polydimethylsiloxane (PDMS) thin film (~200 μm) bonded to glass.

The bottom digital sensor component has femtoliter-sized microwell arrays made by SU-8 molded polydimethylsiloxane (PDMS) thin film (~200 μm) bonded on glass. First, the silicon mold for the microwell structure with a thickness of 4 μm (SU-8 2005, Micro-Chem) was fabricated by photolithography. Second, a precursor of PDMS prepared at a 10:1 base-to-curing agent mass ratio was spin-coated onto the microwell silicon mold (300rpm, 1min). The mold was left on the flat surface overnight and then cured in an oven at 60 °C for 2h. The fully cured PDMS thin film was then transferred to a pre-cleaned 75×38 mm glass substrate through oxygen plasma treatment.

**Surface functioning of the PEdELISA Chip**

The digital sensors are patterned into the microwell arrays at designated locations through a bead patterning process involving several steps. First, a reusable PDMS bead patterning layer, containing long channels running perpendicular to the flow cells' direction, is attached to the microwell array surface. Multiple sets of bead solutions, each targeting a different analyte and with a concentration of 1 mg/mL, are prepared and loaded into separate patterning channels within the bead patterning layer. The beads are allowed to settle inside the microwells for 5 minutes before washing the patterning channels with 200 µL PBS-T (0.1% Tween20) to remove any unbound beads. The bead patterning layer is then removed, and the top PMMA/PSA assembly is permanently bonded to the patterned chip, creating 16 identical flow cells per chip, each capable of multiplexed cytokine measurements. To prevent non-specific binding, SuperBlock$^{TM}$ buffer (ThermoFisher) is loaded into each flow cell. The microarrays are scanned using a microscope to ensure that the bead-filling rate of each array is above 50% and to document the results. Finally, the ports of the chips are sealed with tape, and all chips are stored in a 4°C refrigerator for later assay use.

**Design and fabrication of the PEdELISA fluidic manifold**

A Stereolithography (SLA) 3D printing technique is used to fabricate the microfluidic manifold, which features intricate geometries and multiple internal channels. The manifold is designed with 16 inlets and 8 outlets, each outlet connecting to a rotary valve port on the syringe pump that can be independently activated. The geometry and design of the manifold are optimized to achieve several goals: reducing cross-contamination between samples, balancing throughput and speed, and ensuring a consistent flow profile and volume across 16 flow cells. On the reverse side of the manifold, there are 32 extruded nozzles, each holding an O-ring in place. A CNC machined clamping system was designed to interface with the manifold, creating an airtight seal by sandwiching the PEdELISA chip in the middle (Figure 2C).

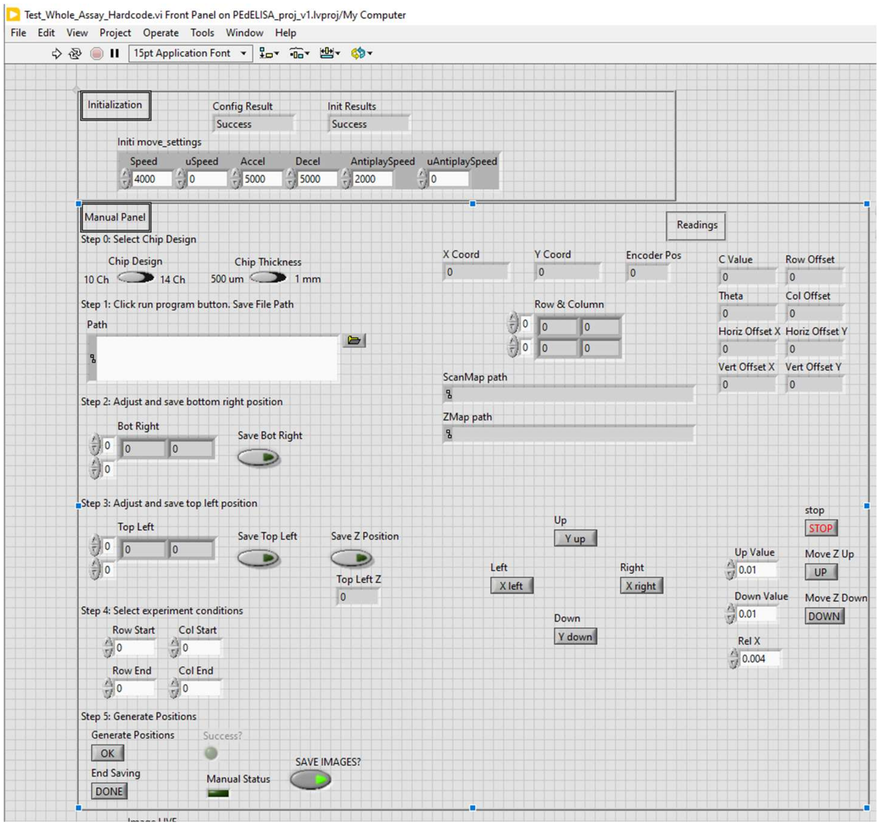


**Figure S2. A user-friendly graphic user interface (GUI) for compact reader control and automatic digital immunoassay data acquisition.**

A compact and cost-effective optical reader system (Figure 2D) was developed for fully automated digital assay imaging, measuring 12" x 12" on a breadboard. The system includes several key components, such as dual-color LED light sources for bright-field and fluorescence imaging, a programmed stepper motorized stage for x,y scanning, and a nanopositioning stage for objective auto-focusing. The fluorescence imaging system is CMOS-based, with imaging quality control provided by a Si photodetector. The system also includes a data acquisition card (DAQ) and a LabVIEW-based user interface and controlling system.

We have developed a user-friendly graphic user interface (GUI) (Figure S2) that enables end-users to perform precise x, y, and z stage movements, specify PEdELISA chip designs, choose a folder path for saving results, and set the starting and ending rows and columns of the microarray on the chip for imaging. Once the user completes the initial setup, loads the PEdELISA chip, and clicks the scanning button, the reader will first perform a quick auto-focus on the microwell structure of the chip and then search for the alignment marks on the upper left and bottom right corner of the chip. We use a machine vision algorithm (Figure S3) for the reader to recognize the alignment marks, detect their position, and use this information to calculate the chip's rotation angle, x and y offset, and eventually obtain the absolute position of all the microwell arrays to be scanned.

Afterwards, the reader will go to 12 pre-defined microarray locations that are evenly distributed on the chip surface, applying a widely adopted auto-focusing algorithm (Sum of Modified-Laplacian (SML)) to obtain the in-focused z value of a stack of images (Figure S3B), use these 12 z values to map the entire chip surface (biharmonic spline, 2D), and interpolate the in-focus z value of every microarray position to be imaged (Figure S3C). Finally, the motorized platform will accurately relocate to the x, y, z coordinates of each microarray, activate the red LED (625nm) to capture brightfield images, and then activate the green LED (554nm) to obtain fluorescence images with exposure time set to be 300ms. This entire process typically takes less than five minutes.

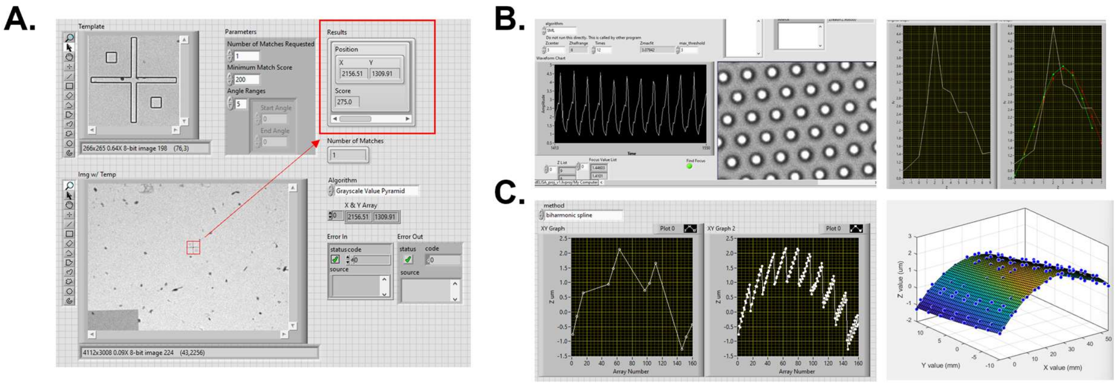


**Figure S3. PEdELISA machine vision algorithm.**

(A) A machine vision algorithm is used to recognize alignment marks, detect their position, and calculate the rotation angle, x and y offsets of a chip. This information is used to obtain the absolute position of all microwell arrays for scanning.

(B) The reader visits 12 fixed microwell array locations evenly distributed on the chip surface. An in-house developed auto-focusing algorithm, Sum of Modified-Laplacian (SML), is used to obtain the in-focus z value at each location.

(C) The 12 z values obtained from the fixed microwell array locations are used to create a map of the entire chip surface using a biharmonic spline in 2D. This map is then used to interpolate the in-focus z value for every microarray position to be imaged.

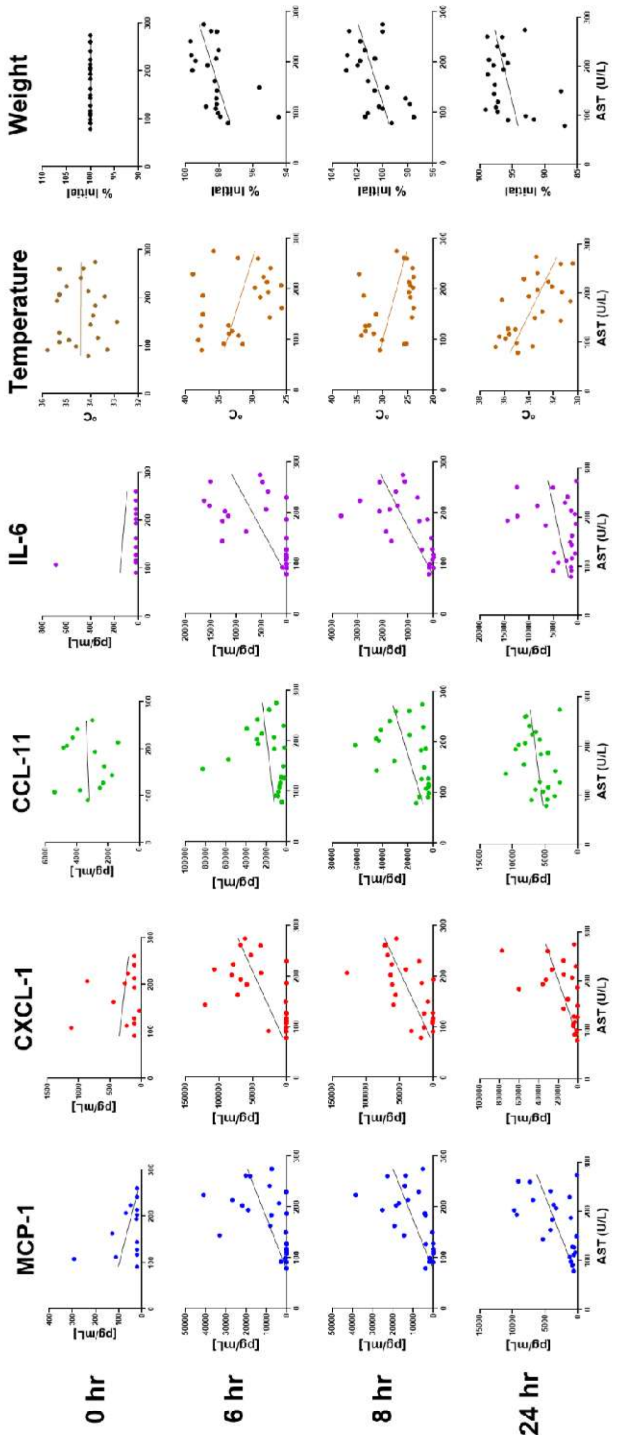


**Figure S4.** Correlation between AST concentration measured at 24 hours post-infection and various parameters: cytokines (MCP-1, CXCL-1, CCL-11, IL-6), body temperature, and percent change in weight (Weight %) at 0, 6, 8, and 24 hours post-injection (rows, across). Weight % was calculated as the percentage change in weight from the baseline measurement at the time of infection (0 hr). Cytokine levels measured at 8 hours post-infection showed the strongest association with liver injury at 24 hours post-infection.

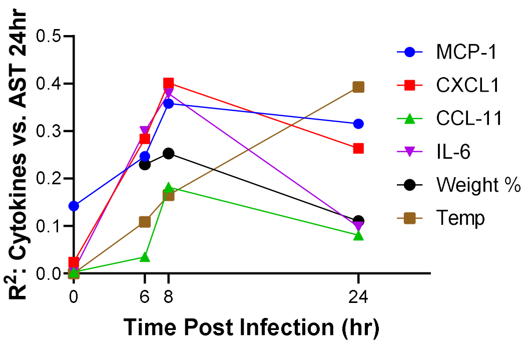


**Figure S5.** $R^2$ values of all cytokines at different times post-infection, correlated with the liver injury marker AST at 24 hours, as measured with the PEdELISA platform, illustrating the importance of time lags in the relationship between immunopathology and organ injury.

| t=2hr | | | 14ul(CS) | | | | | | | | | |
|---|---|---|---|---|---|---|---|---|---|---|---|---|
| **Parameter** | **Degree** | **Scoring** | **tail(0)** | **tail(1)** | **tail(2)** | **tail(3)** | **tail(4)** | **tail(0)** | **tail(1)** | **tail(2)** | **tail(3)** | **tail(4)** |
| Response | Normal response | 4 | 4 | 4 | 4 | 4 | 4 | 4 | | 4 | 4 | |
| to finger poke | Decrease response | 3 | | | | | | | 3 | | | 3 |
| | Severely decreased response | 2 | | | | | | | | | | |
| | Minimal response | 1 | | | | | | | | | | |
| | Noresponse | 0 | | | | | | | | | | |
| Signs of | Normal | 4 | 4 | | | | | | | | | |
| encephalopathy | Tremors,staggering | 3 | | 3 | 3 | 3 | 3 | 3 | 3 | 3 | 3 | 3 |
| | Twisting | 2 | | | | | | | | | | |
| | turning and flipping | 1 | | | | | | | | | | |
| | no response | 0 | | | | | | | | | | |
| | normal | 4 | 4 | 4 | 4 | 4 | 4 | 4 | 4 | 4 | 4 | 4 |
| | piloerecton | -1 | | -1 | | -1 | -1 | -1 | -1 | -1 | -1 | -1 |
| Appearance | periorbital exudates | -1 | | | | | | | | | | |
| | respiratory distress | -1 | | -1 | -1 | -1 | -1 | -1 | -1 | -1 | -1 | -1 |
| | diarrhea | -1 | | | | | | | | | | |
| | | | 12 | 9 | 10 | 9 | 9 | 9 | 8 | 9 | 9 | 8 |

| t=4hr | | | 14ul(CS) | | | | | | | | | |
|---|---|---|---|---|---|---|---|---|---|---|---|---|
| **Parameter** | **Degree** | **Scoring** | **tail(0)** | **tail(1)** | **tail(2)** | **tail(3)** | **tail(4)** | **tail(0)** | **tail(1)** | **tail(2)** | **tail(3)** | **tail(4)** |
| Response | Normal response | 4 | | | | | | | | | | |
| to finger poke | Decrease response | 3 | | | | 3 | | | | | | |
| | Severely decreased response | 2 | 2 | 2 | 2 | | 2 | 2 | 2 | 2 | 2 | 2 |
| | minimal response | 1 | | | | | | | | | | |
| | Noresponse | 0 | | | | | | | | | | |
| Signs of | normal | 4 | | | | | | | | | | |
| encephalopathy | tremors,staggering | 3 | 3 | 3 | 3 | 3 | 3 | 3 | 3 | 3 | 3 | 3 |
| | twisting | 2 | | | | | | | | | | |
| | turning and flipping | 1 | | | | | | | | | | |
| | no response | 0 | | | | | | | | | | |
| | normal | 4 | 4 | 4 | 4 | 4 | 4 | 4 | 4 | 4 | 4 | 4 |
| | piloerecton | -1 | -1 | -1 | -1 | -1 | -1 | -1 | -1 | -1 | -1 | -1 |
| Appearance | periorbital exudates | -1 | | | -1 | | | | | | | |
| | respiratory distress | -1 | -1 | -1 | -1 | -1 | -1 | -1 | -1 | -1 | -1 | -1 |
| | diarrhea | -1 | | | | | | | | | | |
| | | | 7 | 7 | 6 | 8 | 7 | 7 | 7 | 7 | 7 | 7 |

| t=6hr | | | 14ul(CS) | | | | | | | | | |
|---|---|---|---|---|---|---|---|---|---|---|---|---|
| **Parameter** | **Degree** | **Scoring** | **tail(0)** | **tail(1)** | **tail(2)** | **tail(3)** | **tail(4)** | **tail(0)** | **tail(1)** | **tail(2)** | **tail(3)** | **tail(4)** |
| Response | Normal response | 4 | | | | | | | | | | |
| to finger poke | Decrease response | 3 | | | | 3 | | | 3 | 3 | | 3 |
| | Severely decreased response | 2 | 2 | 2 | 2 | | 2 | 2 | | | 2 | |
| | Minimal response | 1 | | | | | | | | | | |
| | Noresponse | 0 | | | | | | | | | | |
| Signs of | Normal | 4 | | | | | | | | | | |
| encephalopathy | Tremors,staggering | 3 | 3 | 3 | 3 | 3 | 3 | 3 | 3 | 3 | 3 | 3 |
| | Twisting | 2 | | | | | | | | | | |
| | turning and flipping | 1 | | | | | | | | | | |
| | no response | 0 | | | | | | | | | | |
| | normal | 4 | 4 | 4 | 4 | 4 | 4 | 4 | 4 | 4 | 4 | 4 |
| | piloerecton | -1 | -1 | -1 | -1 | -1 | -1 | -1 | -1 | -1 | -1 | -1 |
| Appearance | periorbital exudates | -1 | | | | | | | | | | |
| | respiratory distress | -1 | -1 | -1 | -1 | -1 | -1 | -1 | -1 | -1 | -1 | -1 |
| | diarrhea | -1 | | | | | | | | | | |
| | | | 7 | 7 | 7 | 8 | 7 | 7 | 8 | 8 | 7 | 8 |

| t=8hr | | | 14ul(CS) | | | | | | | | | |
|---|---|---|---|---|---|---|---|---|---|---|---|---|
| **Parameter** | **Degree** | **Scoring** | **tail(0)** | **tail(1)** | **tail(2)** | **tail(3)** | **tail(4)** | **tail(0)** | **tail(1)** | **tail(2)** | **tail(3)** | **tail(4)** |
| Response | Normal response | 4 | | | | | | | | | | |
| to finger poke | Decrease response | 3 | | | | | | | | | | |
| | Severely decreased response | 2 | | | | | | | | | | |
| | minimal response | 1 | 1 | 1 | 1 | 1 | 1 | 1 | 1 | 1 | 1 | 1 |
| | Noresponse | 0 | | | | | | | | | | |
| Signs of | normal | 4 | | | | | | | | | | |
| encephalopathy | tremors,staggering | 3 | | | | | | | | | | |
| | twisting | 2 | 2 | 2 | 2 | 2 | 2 | 2 | 2 | 2 | 2 | 2 |
| | turning and flipping | 1 | | | | | | | | | | |
| | no response | 0 | | | | | | | | | | |
| | normal | 4 | 4 | 4 | 4 | 4 | 4 | 4 | 4 | 4 | 4 | 4 |
| | piloerecton | -1 | -1 | -1 | -1 | -1 | -1 | -1 | -1 | -1 | -1 | -1 |
| Appearance | periorbital exudates | -1 | | | -1 | | | | | | | |
| | respiratory distress | -1 | -1 | -1 | -1 | -1 | -1 | -1 | -1 | -1 | -1 | -1 |
| | diarrhea | -1 | | | | | | | | | | |
| | | | 5 | 5 | 4 | 5 | 5 | 5 | 5 | 5 | 5 | 5 |

| t=24hr | | | 14ul(CS) | | | | | | | | | |
|---|---|---|---|---|---|---|---|---|---|---|---|---|
| **Parameter** | **Degree** | **Scoring** | **tail(0)** | **tail(1)** | **tail(2)** | **tail(3)** | **tail(4)** | **tail(0)** | **tail(1)** | **tail(2)** | **tail(3)** | **tail(4)** |
| Response | Normal response | 4 | | | | | | 4 | 4 | 4 | 4 | 4 |
| to finger poke | Decrease response | 3 | | 3 | 3 | 3 | 3 | | | | | |
| | Severely decreased response | 2 | 2 | | | | | | | | | |
| | Minimal response | 1 | | | | | | | | | | |
| | Noresponse | 0 | | | | | | | | | | |
| Signs of | Normal | 4 | 4 | 4 | 4 | 4 | 4 | 4 | 4 | 4 | 4 | 4 |
| encephalopathy | Tremors,staggering | 3 | | | | | | | | | | |
| | Twisting | 2 | | | | | | | | | | |
| | turning and flipping | 1 | | | | | | | | | | |
| | no response | 0 | | | | | | | | | | |
| | normal | 4 | 4 | 4 | 4 | 4 | 4 | 4 | 4 | 4 | 4 | 4 |
| | piloerecton | -1 | -1 | | -1 | -1 | -1 | -1 | -1 | | | |
| Appearance | periorbital exudates | -1 | -1 | | | | | | | | | -1 |
| | respiratory distress | -1 | | -1 | -1 | -1 | -1 | | | | | |
| | diarrhea | -1 | | | | | | -1 | -1 | -1 | -1 | -1 |
| | | | 8 | 10 | 9 | 9 | 9 | 10 | 10 | 11 | 11 | 10 |

**Table S1. Clinical record of the CS mouse model in real-time study.** Clinical evaluation was performed and the scores were recorded at the blood collection time point of 2hr, 4hr, 6hr, 8hr, 24hr.

| Correlation to Cytokine Concentration at 24 hr | | | | |
|---|---|---|---|---|
| Hour | MCP-1 | CXCL-1 | CCL-11 | IL-6 |
| 0 | 0.110 | 0.098 | 0.008 | 0.058 |
| 6 | 0.593 | 0.284 | 0.519 | 0.339 |
| 8 | 0.650 | 0.180 | 0.607 | 0.303 |

**Table S2.** $R^2$ values of cytokines at 0, 6, 8 hours post-infection, correlated with their concentration at 24 hours, as measured with the PEdELISA platform.
